# scACCorDiON: A clustering approach for explainable patient level cell cell communication graph analysis

**DOI:** 10.1101/2024.08.07.606989

**Authors:** James S. Nagai, Michael T. Schaub, Ivan G.Costa

## Abstract

**Motivation:** The combination of single-cell sequencing with ligand-receptor analysis paves the way for the characterization of cell communication events in complex tissues. In particular, directed weighted graphs stand out as a natural representation of cell-cell communication events. However, current computational methods cannot analyze sample-specific cell-cell communication events, as measured in single-cell data produced in large patient cohorts. Cohort-based cell-cell communication analysis presents many challenges, such as the non-linear nature of cell-cell communication and the high variability presented by the patient-specific single-cell RNAseq datasets.

**Results:** Here, we present scACCorDiON (single-cell Analysis of Cell-Cell Communication in Disease clusters using Optimal transport in Directed Networks), an optimal transport algorithm exploring node distances on the Markov Chain as the ground metric between directed weighted graphs. Additionally, we derive a *k*-barycenter algorithm using the Wasserstein-based distance, which is able to cluster directed weighted graphs. We compare our approach with competing methods in several large cohorts of scRNA-seq data. Our results show that scACCorDiON can predict clusters better, matching the disease status of samples. Moreover, we show that barycenters provide a robust and explainable representation of cell cell communication events related to the detected clusters. We also provide a case study of pancreas adenocarcinoma, where scACCorDion detects a sub-cluster of disease samples associated with changes in the tumor microenvironment.

**Availability:** The code of scACCorDiON is available at https://scaccordion.readthedocs.io/en/latest

## 1 Introduction

Single-cell RNA sequencing (scRNA-seq) enables the characterization of cellular processes at unprecedented resolution. Specifically, it allows the study of cell-cell communication (CCC) via the expression patterns of cognate ligand-receptor (LR) pairs across cells detected via scRNA-seq [Armingol et al., 2020, Dimitrov et al., 2022]. As sequencing costs have been reduced by the rapid improvement of single-cell sequencing protocols, it has become possible to create scRNA-seq datasets for large patient cohorts [CZI Single-Cell Biology et al., 2023]. Such datasets, which contain patients in different conditions, hold the potential to improve the understanding of how cell communication changes in various biological settings. However, for a sample-level analysis of such large-scale scRNA-seq patient data, efficient computational approaches are needed [Flores et al., 2023, Joodaki et al., 2024]. For a friendly introduction to the cell-cell communication analysis, we refer to the review written by Armingol et al. [2020, 2024].

However, there are now hundreds of computational methods for LR-based communication analysis [Armingol et al., 2024]. These tools mainly focus on inferring LR pairs within a *single* biological condition. A yet poorly studied aspect is to characterize changes in cell communication in *multiple* biological conditions, such as disease vs. control [Nagai et al., 2021] or over cell differentiation [Li et al., 2022]. To this date, only a few computational methods for CCC – Tensor2Cell and MultiNicheNet – have considered data from multiple samples (patients). MultiNicheNet [Browaeys et al., 2023] builds upon NicheNet [Browaeys et al., 2020], considering both extra-cellular and intra-cellular signaling in CCC. To consider multiple samples, MultiNicheNet obtains pseudo-bulk representations, where cells are bulked for each cell type and sample, and uses a differential expression approach (edgeR, [Robinson et al., 2010]) to perform a multi-conditional differential communication analysis. However, MultiNicheNet is a supervised algorithm that requires the group of samples to be defined prior and does not allow the finding of unknown groups of samples with distinct CCC programs. TensorCell2Cell [Armingol et al., 2022] uses tensor component analysis to detect latent factors explaining changes in CCC associated with sample-level scRNA-seq data. The factors can detect patterns (CCC events) related to individual samples. Similar to MultiNicheNet, Tensor2Cell does not provide any approach for finding unknown groups of samples undergoing distinct CCCs.

In this work, we explore CCC across multiple patients using directed weighted graphs representations. In this representation, cell types are nodes; directed edges represent a communication event connecting a source cell (expressing a ligand) to a target cell (expressing a cognate receptor) [Nagai et al., 2021]. The combined expression of ligand-receptor molecules represents this directed edges’ strength (or edge weight). Using a graph representation enables us to exploit a large variety of graph algorithms, such as pagerank [Page et al., 1998], to detect latent cell-cell communication events leading to fibrosis [Jansen et al., 2022, Leimkühler et al., 2020, Peisker et al., 2022]. Within the sample-level cell-cell communication context, a common challenge in the sample-level analysis is clustering the samples. When using a graph-based representation, this corresponds to clustering a set of graphs according to their similarity, which is a computationally hard task. This problem has been previously tackled with graph-based optimal transport approaches, which (implicitly) utilize spectral properties of graphs [Maretic et al., 2022, Petric Maretic et al., 2019] or node distances [Chowdhury and Mémoli, 2019, Scholkemper et al., 2024, Xu et al., 2019]. However, these approaches can only be used in undirected graphs and would miss important information regarding the directionality of LR interactions.

## 2 Approach

We propose scACCorDiON (single-cell Analysis of Cell-Cell Communication in Disease clusters using Optimal transport in directed Networks). scACCorDiON represents the CCC data of each sample as a directed weighted graph (DWG) and employs an optimal transport (OT) to compute the Wasserstein distance [Bonneel et al., 2011] between CCC graphs. For this, we assume that the probability masses to be transported via OT correspond to the strength of the LR communication pairs (i.e., expression values). To model this, we lift each CCC graph to a line graph, where a node represents a directed interaction, and its mass (or weight) represents the LR expression values. To define a cost (distance) function for the transport between the nodes of this line graph, we use a Markov Chain based distance measure [Boyd et al., 2021], which can be used as a cost function with OT in directed weighted graphs [Nagai et al., 2024].

Moreover, we leverage the fact that OT enables us to estimate barycenters of a set of CCC networks [Cuturi and Doucet, 2014]. Barycenters represent a group of CCC graphs and can be used together with transport maps for interpretation, i.e., delineating cell communication events that change between groups of samples. Specifically, these barycenters can used as a “centroid” values within an expectation-maximization clustering algorithm denoted *k*-barycenters.

We benchmarked scACCorDiON and baseline methods (i.e., methods based on the tabular representation of the data and/or ignoring graph structures) on six large scRNA-seq cohorts with up to 126 samples and up to millions of cells. We did not benchmark MultiNicheNet or Tensor2Cell, as neither method explicitly provided methods to find sub-groups of samples.

We evaluate the performance of clustering in recovering known labels of the samples. Finally, we explore CCC barycenters and transport maps to characterize CCC events on a pancreas adenocarcinoma scRNA-seq data [Peng et al., 2019] and find that scACCorDiON can detect so far uncharacterized sub-groups of disease samples and it’s main communication shifts. We also compared these results with latent spaces derived from Tensor-cell2cell, and factors were able to depict differences across the new clusters. Although interactions selected by the factors corroborate with interactions depicted by scACCorDiON, a mapping between interactions was not obtained. This indicates scACCorDiON’s unique capability to group and interpret disease-related cell-cell communication programs featuring interaction shifts between disease sub-stages.

## 3 Material and Methods

### 3.1 scAccordion

scACCorDiON is an optimal transport-based framework for directed weighted graph metric learning and clustering. The input to scACCorDiON is a set of directed weighted CCC graph {𝒢^1^, …, 𝒢 ^*p*^} containing *p* graphs. We assume that each of these *p* graphs has been obtained from a ligand-receptor analysis method [Dimitrov et al., 2022, Nagai et al., 2021], applied to a scRNA-seq dataset containing multiple samples (e.g., a cohort). The CCC graph of sample *k* is then defined as 𝒢^*k*^ = (*V*^*k*^, *E*^*k*^, *w*^*k*^), where a node *v* ∈ *V*^*k*^ represents a cell type, and a directed edge *e* ∈ *E*^*k*^ connects a pair of cell types when these cells are predicted to be communicating through a ligand (source cell) and receptor (target cell) pair. The weights of the edges *w*^*k*^ are related to the amount of communication between the source and target cell, e.g., *w*^*k*^(*e*) is the sum of all ligand-receptor expressions (LRScore). Note that in our problem setup, nodes can be identified across all graphs for a given scRNA-seq dataset, i.e., all samples *k* have the same cell types and thus the same node-set *V*^*k*^ = *V*.

#### 3.1.1 Metric Learning From Directed Weighted Graphs

scACCorDiON uses OT to obtain a metric between patients’ CCC graphs [Bonneel et al., 2011]. More specifically, our hypothesis is that the edge weights of each graph are a specific realization of a signal supported on the same underlying graph structure, i.e., every CCC graph has distinct signals related to the LR expression between cell pairs, but they all share the same topology. Therefore, the directed weighted graph OT (DW-OT) problem we consider here consists of finding an OT map between the CCC graph edge signals with respect to an edge-to-edge cost (distance) matrix C. For this, we proceed in two steps. First, we define a shared topology line graph, which encodes how LR communication signals feed into each other across all samples. Second, we treat each sample’s edge weights as a signal distribution on this line graph and use OT to compare two such distributions. For the latter step, we also need to define a distance matrix C for the nodes in the (directed) line graph, which we do by borrowing ideas from Markov chain theory [Young et al., 2015a,b].

##### Shared Topology Graph (STG)

We build a *directed line graph* 𝕃 = (𝕍, 𝔼, 𝕎), whose vertex set 𝕍 contains each possible edge (*j, k*) contained in one of the sample graphs 𝒢^*i*^. The edge set 𝔼 of the linegraph is defined as follows: an edge *e*^′^ = (*u*^′^, *v*^′^) ∈ E exists if the target of *u*^′^ is the source of *v*^′^, i.e., the target node of edge *u*^′^ in the original graph, is the source node of edge *v*^′^ in the original graph. Note that this line graph’s structure essentially encodes the union of interactions of all CCC graphs. We denote this graph as the “shared topology graph” (STG). Finally, we defined the weight *w*_*u*_^′^*v*^′^ of edge (*u*^′^, *v*^′^) as the proportion of graphs containing both edges *u*^′^ to *v*^′^. This makes transport of masses between common edges (in the original graphs) more likely than transport between rare edges (in the original graphs). As our line graph can have unconnected components, in general, we add to the STG a low-rank regularization term, as popularized within the context of the well-known PageRank algorithm [Gleich, 2015, Page et al., 1998], to obtain a well-posed problem. Specifically, this guarantees the global reachability of all nodes in the STG, which is required to compute the distance between nodes in a graph. Using the STG, we can represent each sample as a signal distribution on the nodes of the STG, which we can compare via OT. However, for the computation of OT, we also need to define a distance matrix for the nodes on the STG, which specifies the cost of moving a signal from one node to another node in the STG.

##### Hitting Time Distance (HTD)

Here, we consider the Hitting Time Distance (HTD [Boyd et al., 2021]), which is a metric that can be applied to *directed* weighted graphs. To derive the HTD, consider a discrete-time Markov chain (*X*_*t*_)_*t*≥0_ defined over the vertices 𝒱 = {1, …, *N*} of a strongly connected graph. We assume the chain has a starting distribution λ and an irreducible transition matrix **P** = **D**^−1^**A**, where *A* is the adjacency matrix of the shared topology graph and **D** = diag(*A***1**). The Markov chain can then be described according to the state transition probabilities:

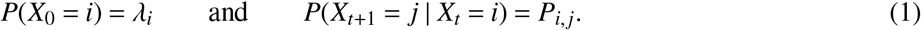

Let π ∈ ℝ^*N*^ be the invariant distribution of the chain, i.e., π**P** = π. For a starting point distributed according to λ, the *hitting time* of a vertex *i* ∈ 𝒱 is the random variable τ_*i*_ = inf{*t* ≥ 1 : *X*_*t*_ = *i*}. Following [Boyd et al., 2021], we define the probability that starting in a node *i*, the hitting time of *j* is less than the time it takes to return back to *i* by *Q*_*i, j*_ := *P*(τ _*j*_ ≤ τ_*i*_ | *X*_0_ = *i*). Based on the matrix **Q** = [*Q*_*i, j*_] a normalized hitting time matrix **T** can be defined in terms of its entries

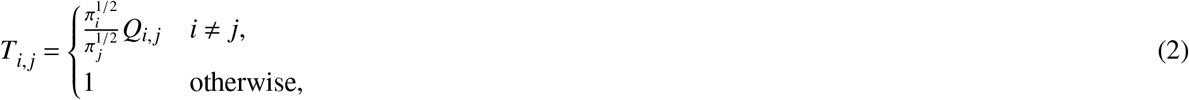

If **P** is an irreducible stochastic matrix, i.e., the underlying graph is strongly connected, the Hitting Time Distance Matrix can be obtained by:

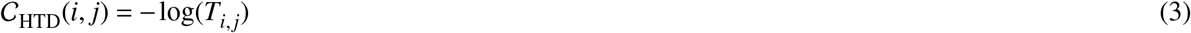

This distance can now be used as cost matrix *C* for an OT problem that considers the movement of signal masses on the STG for different samples.

##### Computing a graph-based CCC distance

To set up our OT-based distance, let us collect the edge weights of each CCC graph in a matrix ℙ ∈ ℝ^*p*×*E*^, where *E* is the size of the union of the edge sets of all graphs (samples). Stated differently, *E* corresponds to the number of nodes in the line-graph. Hence, the columns of ℙ are indexed by the (directed) edges and rows by the samples/graphs, i.e., the row ℙ_*k*,:_ describes the edge-weights *w*^*k*^ of the *k*th graph 𝒢^*k*^, which is appropriately zero padded, in case 𝒢^*k*^ does not contain certain edges which are present in other graphs.

The optimal transport map Γ^∗^ ∈ ℝ^*E*×*E*^ for two probability distributions defined on the nodes of the line graph as induced by the two CCC graphs 𝒢^*k*^ and 𝒢^*l*^ can now be computed as

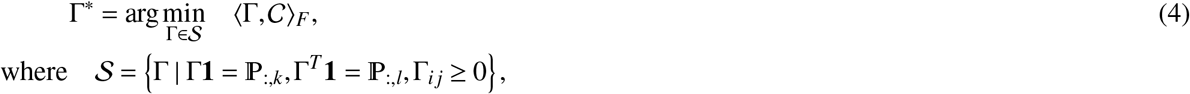

and the associated (induced) Wasserstein distance between the two CCC samples is:

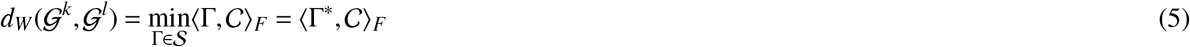

#### 3.1.2 Clustering Patient’s Networks

scACCorDiON employs the metric *d*_*W*_ (Eq. 5) to perform clustering of CCC graphs. One approach for this is a *K*-medoids partitioning algorithm [Rdusseeun and Kaufman, 1987, Schubert and Rousseeuw, 2019, 2021], which only requires a distance matrix and detects samples (medoids) as representative of clusters. scACCorDiON leverages the fact that we can also compute barycenters for distributions of (directed, weighted) graphs via the Wasserstein optimal transport framework [Cuturi and Doucet, 2014].

**K-barycenters clustering** A Wasserstein barycenter of a set of graphs **G** = {𝒢^1^, …, 𝒢^*p*^} can be defined as:

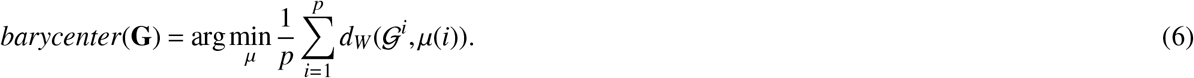

Time-efficient solutions to this problem can be obtained by using a dissimilarity-based loss function (Sinkhorn) of the optimal transport algorithm [Cuturi and Doucet, 2014]. In addition, we use barycenters to define an expectation-maximization-based clustering algorithm, where barycenters represent the “centroids” and we use the Wasserstein distance (Eq. 5) between graphs and barycenters. Given *k* as the number of desired clusters, *Y* be an indicator variable, where *y*_*i*_ ∈ {1, …*k*} indicates the cluster of 𝒢^*i*^, and {μ^1^, …, μ^*k*^} indicates the set of barycenters, this leads to clustering algorithm(Algorithm 1).

##### Algorithm 1 K-Barycenters

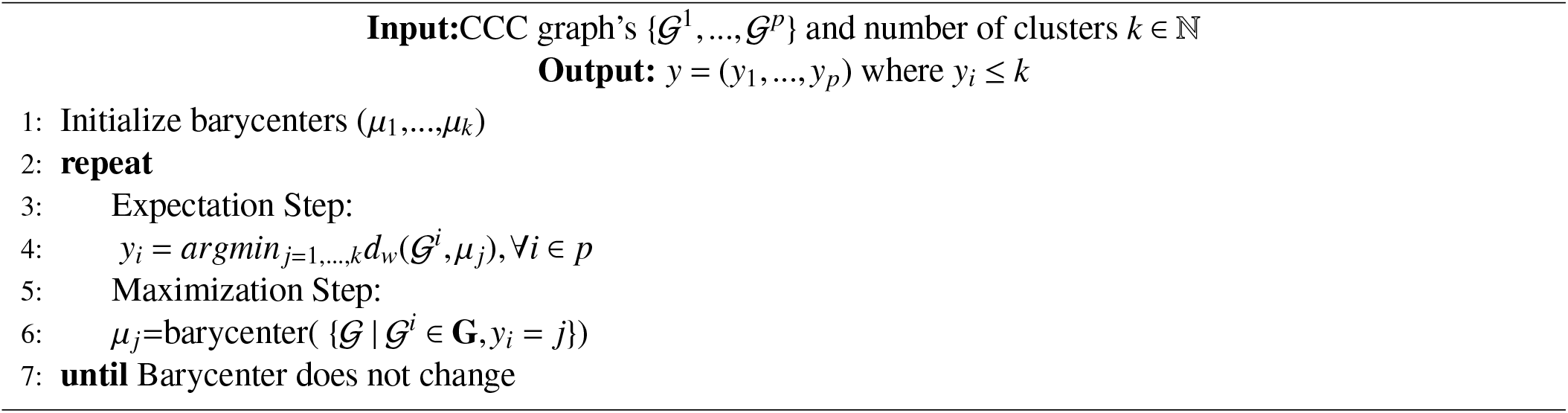

To avoid local maxima due to the random initialization, we repeat the optimization process 100 times and select the solution with the lowest average Wasserstein loss per cluster. Moreover, we use a seeding process to pick the initial barycenters based on selecting CCC graphs that maximize the cluster-to-cluster distance as described in Arthur et al. [2007].

### 3.2 Benchmarking

For benchmarking, we have collected six publicly available disease scRNA-seq cohorts, from which samples were annotated with their disease status. We obtained pre-processed, integrated, clustered, and annotated objects for all data sets from [CZI Single-Cell Biology et al., 2023]. An exception is the pancreas adenocarcinoma data sets, which were pre-processed as described in [Joodaki et al., 2024].

The LR inference was performed with CellPhoneDB [Efremova et al., 2020] implemented in LIANA [Dimitrov et al., 2022] framework by only considering cells in a patient sample. The parameter related to the minimum expression proportion for the ligands/receptors is set to *exp*_*p*_*rop* = 0.15, and highly significant interactions were considered *p* − *value* ≤ 0.01. A description of the data set’s main features is provided in Table 1. While scACCorDiON can be used with any LR inference algorithm, we choose CellPhoneDB due to its widespread use in the literature. We also note that CellPhoneDB is the best-performing tool in the recent single-cell benchmark. [Luecken et al., 2024]. CCC graphs were generated using CrossTalkeR [Nagai et al., 2021].

**Table 1.**
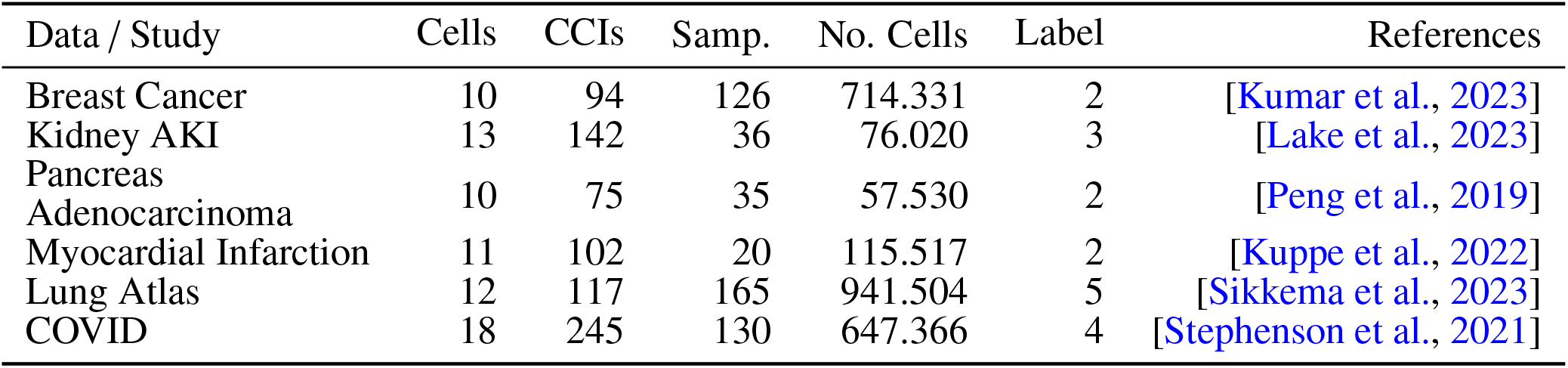
Main features of data sets used in the benchmark, including the number of cell types (Cells), the average number of directed cell-cell interactions (CCI) detected with ligand-receptor analysis, number of individuals/samples, number of cells, and number of sample labels.

We are unaware of another computational approach that can cluster samples by considering CCC information. However, we can contrast scACCorDiON with the following baseline approaches. First, we consider tabular representations (edge weight matrix lP) of the data as input (Tabular. Due to the high dimensionality of lP, we first perform a dimension reduction with Principal Component Analysis (PCA). As a OT baseline method, we compute the correlation distance on the matrix lP as a cost function for the previously described OT framework. This approach is denoted CORR-OT. Note that the last approach does not consider the graph’s topology.

For distance metrics obtained by evaluated methods (Tabular, CORR-OT and DW-OT, we run either *k*-barycenters, *k*-medoids and the community detection algorithm Leiden [Traag et al., 2019]. The last algorithm is chosen based on its wide-spread use in scRNA-seq pipelines [Wolf et al., 2018]. Note also that for the for Tabular, we use *k*-means algorithm as this is equivalent to a *k*-barycenter in an Euclidean space. Algorithms were run by varying the number of clusters from 2 to 7. The Adjusted Rand Index [Hubert and Arabie, 1985](ARI) and Rand Index (Rand) for the *k* equal to the number of class labels and *k* with maximum ARI value was computed for each clustering. Here, the disease labels are used as true classes. Friedman-Nemenyi test for every metric, clustering, and distance combination to address statistically the rank differences [Demšar, 2006, Nemenyi, 1963]. For Leiden [Traag et al., 2019], we vary the resolution parameter to 0 and 1 with 0.01 steps, as this allows obtaining distinct clusters.We refer to Supp. Fig. 1 for an overview of the experimental design. To explore the interpretation ability of Tensor-cell2cell [Armingol et al., 2022], we also estimated tensor decomposition and contrasted results with the disease labels and the new clustering by scACCorDiON. Here, we followed the tutorial available in https://liana-py.readthedocs.io/en/latest/notebooks/liana_c2c.html. Elbow optimization was performed to selected the optimal number of factors (8).

## 4 Results

### 4.1 Benchmarking cell-cell communication graph clustering

We evaluated the performance of scACCorDiON and baseline competing methods using six publicly available scRNA-seq cohort datasets. The datasets contain between 10 and 33 cell types, 20 and 165 samples, and 57,530 to 941,504 single cells. CCC graphs have an average interaction number between 75 and 142 (Table 1). scACCorDiON’s mainly consists of using a *k*-barycenter clustering algorithm with a Wasserstein distance considering both the direction and topology of graphs (DW-OT). In the evaluation, we include a baseline OT method (CORR-OT), which considers the signal directions but not the topology of the CCC graphs. In addition, we include a simple approach using the graph signal matrix lP Tabular as input, which is given as input to *k*-means (note that k-means is equivalent to the *k*-barycenter in the case of data points). To evaluate the impact of the clustering method, we also performed clustering for all methods with *k*-medoids and Leiden algorithm [Traag et al., 2019]. All methods are evaluated regarding their performance in finding clusters associated with the known class labels of samples as measured by the ARI [Hubert and Arabie, 1985]. Class labels indicate the individual’s health status: healthy vs. diseased (or disease sub-type). ARI is measured for the number of *k* equal to the number of true labels or the maximum ARI after varying the number of *k* from 2 to 7 (or cluster resolution) for a given data set and algorithm. The corresponding individual line plots are displayed in Sup. Fig. 2. Additionally, we repeated the evaluation assay using the rand index [Rand, 1971], considering the agreement of two partitions without any correction.

Benchmarking results are displayed in Fig. 2A-D. We observe that DW-OT with k-barycenter has the highest mean ARI for the maximum ARI evaluation (Fig. 2A). A Friedman-Neymeni test indicates that DW-OT with *k*-barycenter has the highest ranking and significantly outperforms Tabular based baseline approaches and the *k*-medoids CORR-OT (Fig. 2B). DW-OT with *k*-medoids obtains the highest mean ARI for the number of clusters equal to the number of classes (Fig. 2C-D). A Friedman-Neymeni test indicates that also this *k*-medoids variant outperforms PCA and CORR-OT with a *k*-medoids algorithm. Fig. 2E shows embeddings [Moon et al., 2017] obtained from distances generated in this study. Regarding Rand Index, we observe similar rankings of methods and an average dRand index varying from 0.6 to 0.8 for DW-OT. These results underline the advantage of DW-OT, which is the only approach incorporating both directionality and connectivity of CCC graphs to cluster the samples. Of note, the Leiden algorithm [Traag et al., 2019], commonly used in single-cell analysis, is outperformed by *k*-medoids and *k*-barycenters. A potential explanation is that the algorithm depends on a high number of samples, which is not present in sample leven analysis.

While we cannot observe a significant difference between the clustering performance of *k*-medoid and *k*-barycenter clustering with DW-GOT, another essential aspect is the explainability of the clustering representatives. Therefore, we compare the signals of barycenters and medoids in the myocardial infarction data. Of note, both algorithms reach the same clustering solution in the myocardial infarction data and detect a cluster only containing ischemic samples (Fig. 3A). A comparison of the signals of the barycenter and medoids for the ischemic group indicates that for most interactions, medoids tend to have values with higher deviation from the barycenter (Fig.3B). The furthest points of the diagonal on Fig.3B include Fibroblast-to-Fibroblast to endothelial cells (Fib− >Fib/Endo) and Proliferating cells to Proliferating cells or Endothelial cells (Prolif− >Proli/Endo). For these cell-cell interactions, we observe that the medoid has a higher error in representing signals of CCC graphs in the same cluster than the barycenter (Fig. 3C). Altogether, these results indicate that medoid signals are more prone to the presence of outlier values than barycenters.

**Figure 1.**
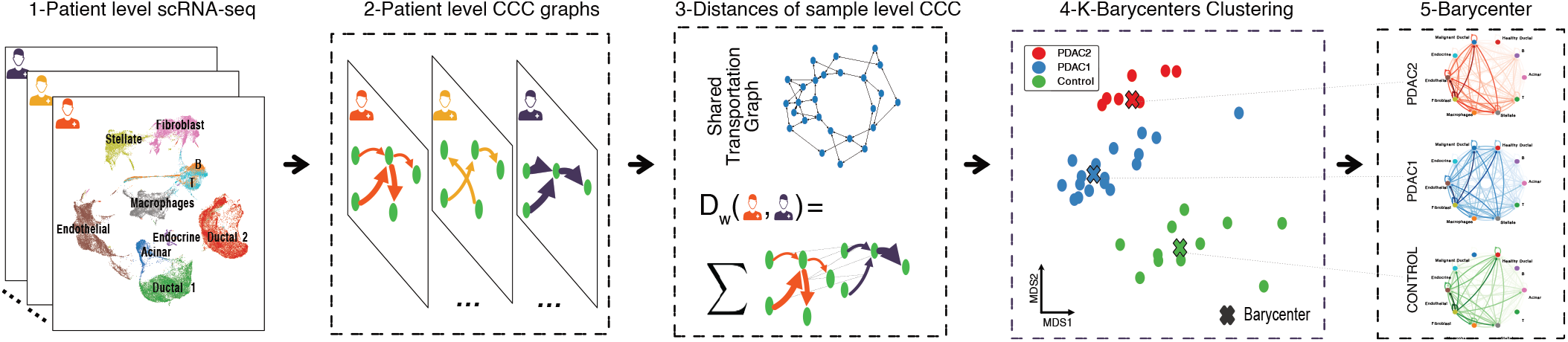
Overview of scACCorDiON: scACCorDiON receives scRNA-seq cohort experiment as input. In the first step, ligand-receptor analysis is performed, and the cell-cell communication graphs for every sample (patient) are recovered. Second, we use graph-based optimal transport, which considers directions and weights of cell-cell interactions, to measure the distance between all pairs of samples. In the third step, this distance is used as input a) for *k*-medoids and *k*-barycenter algorithms that find groups of CCC graphs and produce representative CCC graphs for different patient conditions, and b) for dimension reduction algorithms to create low-dimensional data visualizations. In summary, scACCorDiON enables the analysis of patient cohorts at a CCC graph level and allows for a quantitative comparison of the changes in CCC graphs using optimal transport. Moreover, Barycenters/Medoids are a proxy to facilitate the explainability of the produced results.

**Figure 2.**
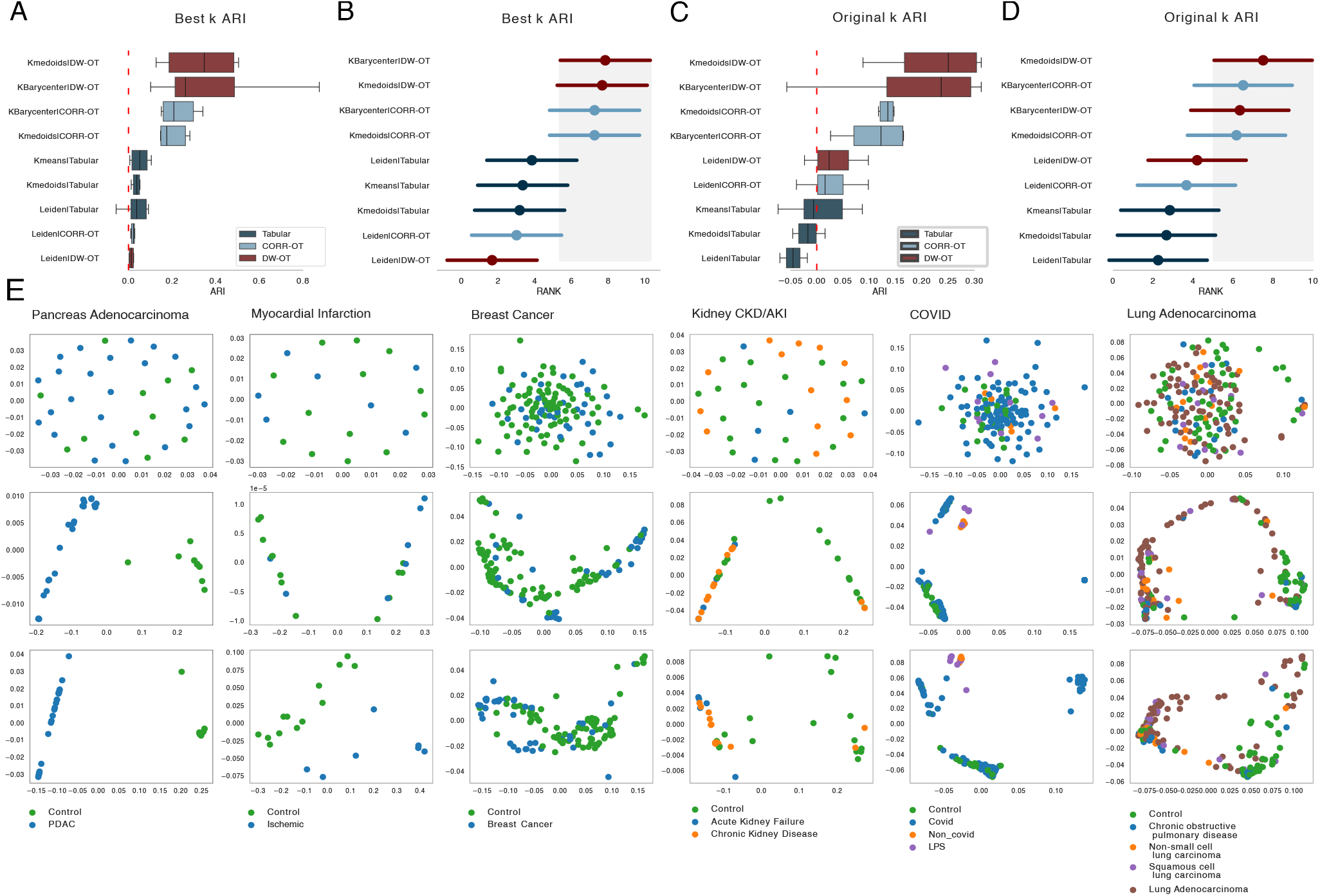
Clustering Benchmark: A) Boxplots indicate the maximum ARI value distribution (x-axis) distribution for all evaluated methods over five scRNA-seq data sets. B) Ranking values (mean and std) for each method and data set regarding the maximum ARI value. The highest ranking indicates the highest ARI. The gray area indicates the 95% confidence interval of the Friedman and Nemenyi posthoc-test). Methods whose average values are not within the gray area have significantly lower rankings than the top-ranked methods. For both A and B, methods are ranked by average in decreasing order. C and D is the same as A and B for the ARI estimates, with the number of clusters equal to the number of labels. E) PHATE two-dimensional embeddings of the distances matrices estimated by Tabular, CORR-OT, and DW-OT for all evaluated data sets. Colors correspond to true labels.

**Figure 3.**
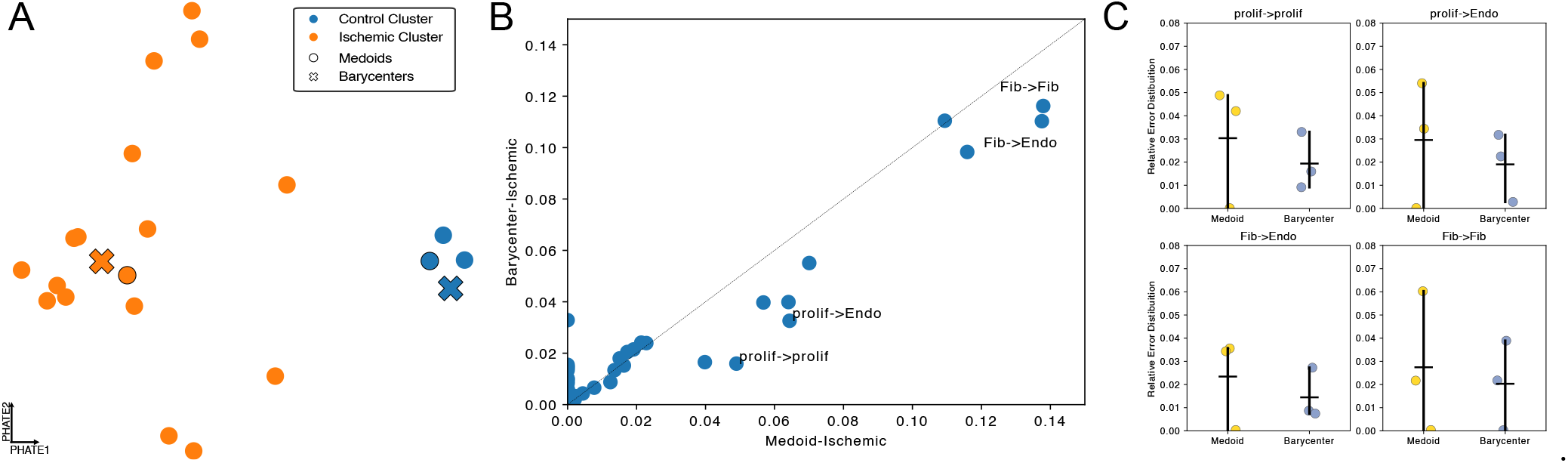
Comparison of Medoids and Barycenters: A) PHATE embedding with clustering results and representatives (medoids and barycenter) in the myocardial infarction single-cell data. B) A scatter plot comparing the cell-cell communication signals between medoids (x-axis) and barycenter (y-axis) for the ischemic cluster. Points farther from the diagonal are highlighted. The shorthand Fib. denotes fibroblasts, endo. denotes endothelial cells and prolif. stands for proliferating cells. C) Difference of signal of barycenter and medoids signals (y-axis) with all samples of the ischemic cluster for selected cell-cell interactions.

### 4.2 scACCorDiON detects a sub-cluster of Pancreas Adenocarcinoma

To evaluate the power of scACCorDiON in the detection of novel sub-clusters, we perform a Silhouette anal-ysis [Rousseeuw, 1987] to identify data sets with a higher number of clusters than true labels (Supp. Fig 2). Interestingly, we observe that for the pancreas adenocarcinoma (PDAC) data, scACCorDiON predicts a sub-cluster associated with controls and two sub-groups associated with PDAC samples (Fig. 4A). As displayed in Fig. 4B, PDAC 1 has overall increased communication, particularly interactions related to ductal, malignant ductal, and fibroblast cells. PDAC 2 demonstrates a loss of communication regarding Acinar and Ductal cells, while we observe an increase in communication-related to Malignant Ductal, B, and Endocrine cells. The prominent signal related to the Malignant Ductal Cells indicates that PDAC 2 clusters are, possibly, linked to more advanced disease stages than PDAC 1 cluster [Peng et al., 2019].

**Figure 4.**
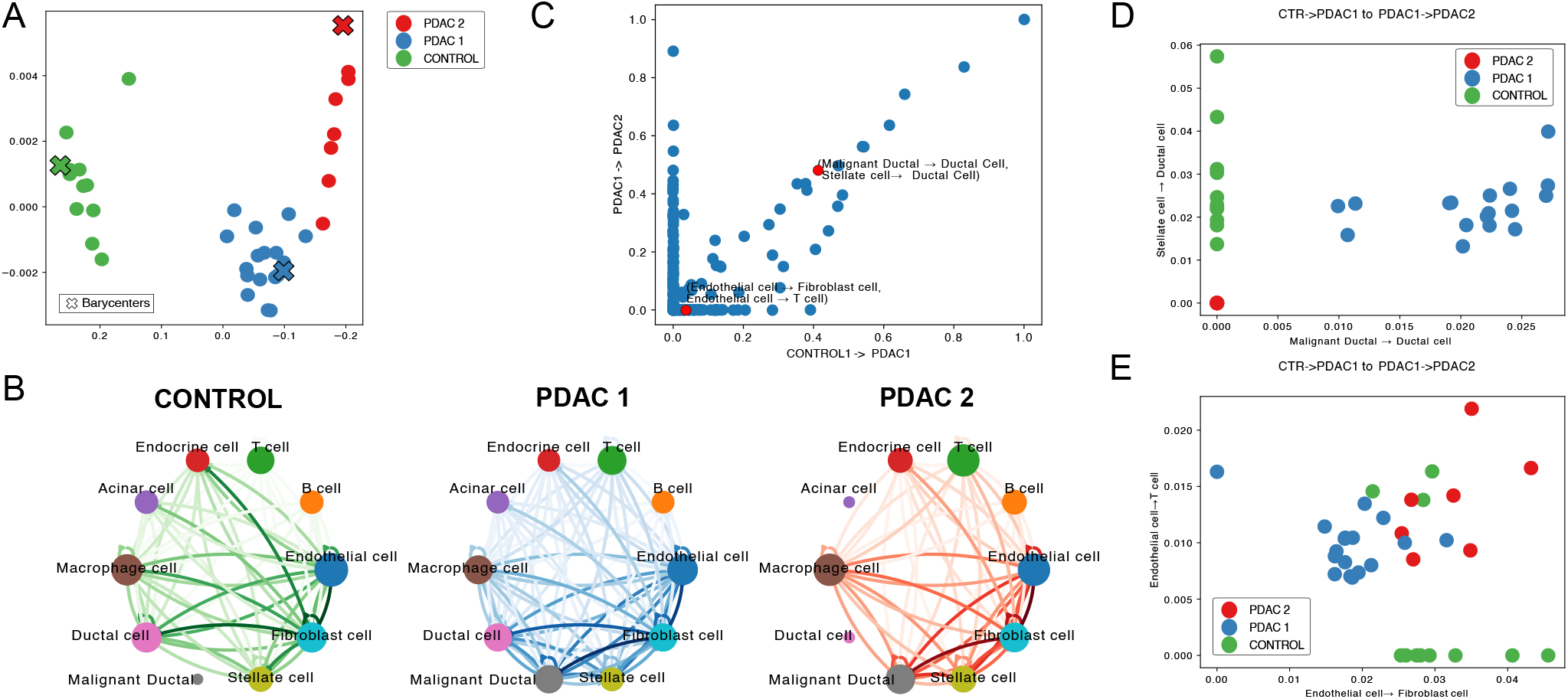
Sub-cluster analysis in pancreas adenocarcinoma. A) MDS plot containing the clustering results on pancreas adenocarcinoma. One cluster contains all control samples (Control), and two clusters contain disease samples (PDAC 1 and 2). B) CCC graphs of barycenters of the detected sub-clusters. Edge thickness indicates the strength of the cell-cell interactions. Node sizes are placed accordingly to the node pagerank C) Scatter plot with the transport map between the barycenter of the control and PDAC 1 (y-axis) and PDAC 1 and PDAC2 clusters (y-axis). Every dot shows the mass transported between two cell-cell pairs in the comparisons. D-E) Scatter plot with signals associated with selected cell-cell pairs for all the samples shown in A). Colors correspond to the sample cluster.

To understand CCC events related to transitions from control to early disease (Control -> PDAC 1) and between mild and advanced disease (PDAC 1->PDAC 2), we contrast the transport maps (Γ) between the barycenters of these pairs of groups (Fig. 4C). We observe that pairs of CCC interactions with high transport masses discriminate well the detected groups (Fig. 4D-E). For example, one of the highest transport masses, observed in Fig.4D, is associated to the map between the interaction (Stellate cells and Ductal cells) and (Malignant Ductal and Ductal cell) which may indicate an triggering event in the realm of Malignant Ductal cells and a compleate transition of Ductal cells to Malignant Ductal cells in the PDAC2 stage. In Fig.4D, we observe a loss in the communication of Endothelial cells towards Fibroblast cells in CONTROLS to an increase in the communication of Endothelial cells towards T cells. This indicates a possible mechanism of immune response occurring upon injury in Pancreatic tissue.

We next make use of the LR analysis from CrossTalkeR [Nagai et al., 2021] to further investigate the interactions associated with the communication between Malignant Ductal cells and Ductal cells (Supp. Fig. 4). We observed high expression in ERBB and EGFR receptors’ interactions among the top LR pairs. These receptors were previously assigned to be related to pancreatic intraepithelial neoplasia (PanIN) [Ghasemi et al., 2014, Meyers et al., 2020], described as a precursor stage of Pancreas Adenocarcinoma. Moreover, ligands secreted by Malignant Ductal cells include matrix metalloproteinase-7 (MMP7) and galactins (LGALS-3/LGALS-3BP/LGALS-9), which have been recently shown to be expressed in malignant cells [Crawford et al., 2002]. These results support an association between CCC changes in the tumor microenvironment and PDAC progression.

As an alternative to the previous analysis, we also explore the use of Tensor-cell2cell [Armingol et al., 2022], which allows a factor-based and sample-level interpretation. If we check the factors by comparing the two known class labels (Control vs. PDAC), we observe that two factors (6 and 8) are significantly associated with controls; 5 factors (2, 3, 4, 5 and 7) with PDAC and one factor (1) is not related to the known labels (Supp. Fig. 5A). Interestingly, by providing the clustering from scACCorDiON, we observe that Factor 1, which was not discriminative between Controls and PDAC groups, is related to PDAC2 samples. In summary, we have Factors 1 and 7 related to interactions related to PDAC2. Factors 2,3,4 and 5 show interactions related to PDAC1. Lastly, Factors 6 and 8 are related to control samples (Supp. Fig. 5B). This supports the biological relevance of these clusters.

For interpretation, Tensor-cell2cell allows the estimation of cell-cell networks related to each factor. Factor 1 (PDAC 1 high) highlights, for example, the interaction from Malignant Ductal cells, including interactions towards Ductal cells. While this finding is on par with our the previous analysis, Tensor-cell2cell do not provide transport maps between the groups, which could explain changes in communication related to the disease subtypes as revealed by scACCorDiON.

## 5 Discussions and Conclusion

Single cell-based LR analysis enables the inference of CCC events related to complex diseases. However, sample-specific analysis, crucial for understanding CCC events in patient cohorts, has only been addressed to a limited extent so far. Here, we explore the problem of clustering samples that share similar cell communication patterns by modeling sample-specific CCC as directed and weighted graphs. We propose a graph-based optimal transport framework that finds optimal probabilistic mappings between cell communication signals and cell cell graphs. Moreover, this framework allows us to measure the distance between any two directed graphs (regarding a Wasserstein distance) and estimate “average directed weighted graphs” (barycenters) representing typical CCC patterns within a group of samples. Our algorithm is currently unique in that it allows both computing distances and clustering of directed weighted graphs.

We have applied our DW-OT algorithm to calculate CCC graphs estimated in scRNA-seq with large cohorts and found that it outperforms other algorithms. We further showcased how both barycenter and transport matrices can be used to interpret communication events supporting detected clusters. This is exemplified in the pancreas adenocarcinoma data set, where DW-OT detected sub-clusters not characterized in the original study presenting the data [Peng et al., 2019].

More importantly, the detected events reveal tumor microenvironmental changes potentially related to disease progression. We also showed that combining scACCorDiON groups with factors from Tensor-cell2cell allowed an interpretation of the main cell-cell interactions driving distinct groups. However, using Tensor-cell2cell does not provide a representative of each cluster nor allows a detailed understanding of possible interaction shifts as provided by scACCorDiON. Future challenges include extending the DW-OT framework to work at the LR level. This would require algorithms dealing with potentially large and noisy LR networks. We also noted that batch effects, frequently present in scRNA-seq cohort data, can affect sample level as analysis as well as of scRNA-seq data [Joodaki et al., 2024]. This opens a new venue for improvements in the current OT methods.

## Data availability

Code is available at https://github.com/jsnagai/scACCorDiON/ and https://scaccordion.readthedocs. io/en/latest/. The data and results are available upon request at https://zenodo.org/doi/10.5281/zenodo.10808382

## Supplement

**Supp Fig 1.**
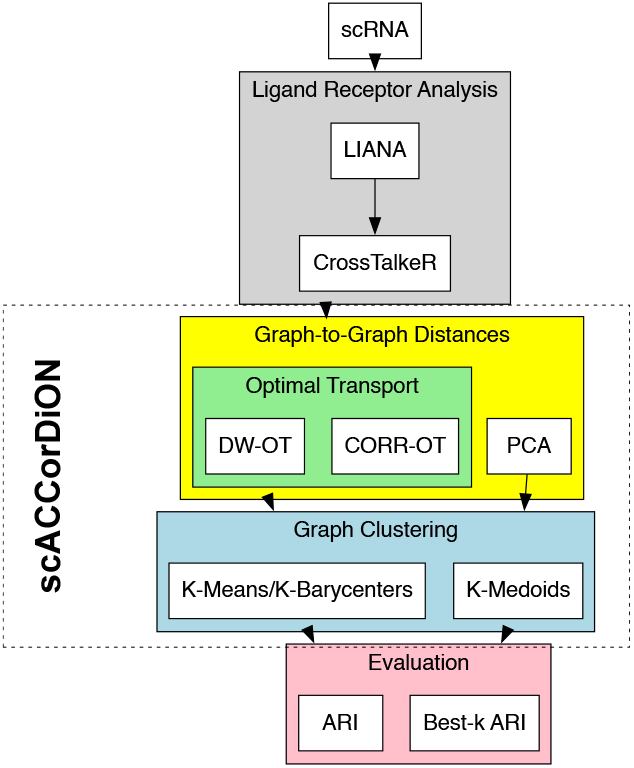
Flowchart with the experimental design of this study and an overview of scACCorDiON. From the input scRNA cohort to the clustering benchmark

**Supp Fig 2.**
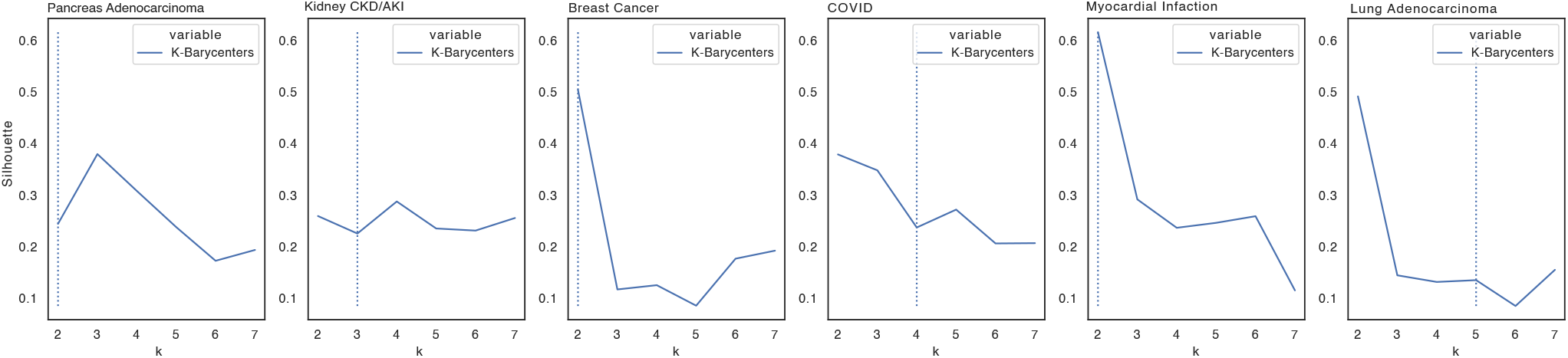
Silhouette of different clustering methods using the DW-OT. Traced lines indicate the number of labels associated with each data set.

**Supp Fig 3.**
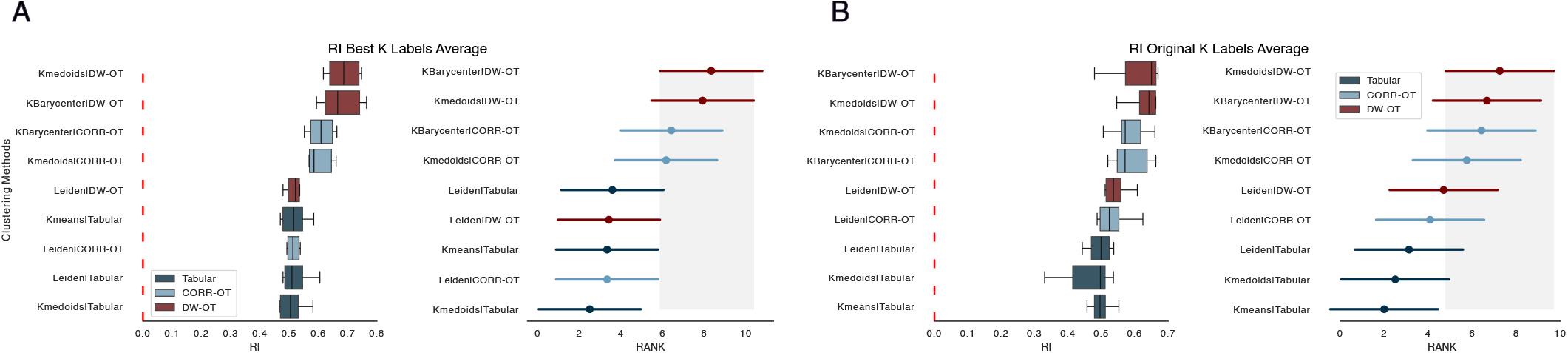
A)Boxplots indicate the maximum Rand Index(RI) value distribution (x-axis) distribution for all evaluated methods over five scRNA-seq data sets. B) Display the ranking values (mean and std) for each method and data set regarding the maximum RI value. The highest ranking indicates the highest RI. The gray area indicates the 95% confidence interval of the Friedman and Nemenyi posthoc-test). Methods whose average values are not within the gray area have significantly lower rankings than the top-ranked methods. For both A and B, methods are ranked by average in decreasing order

**Supp Fig 4.**
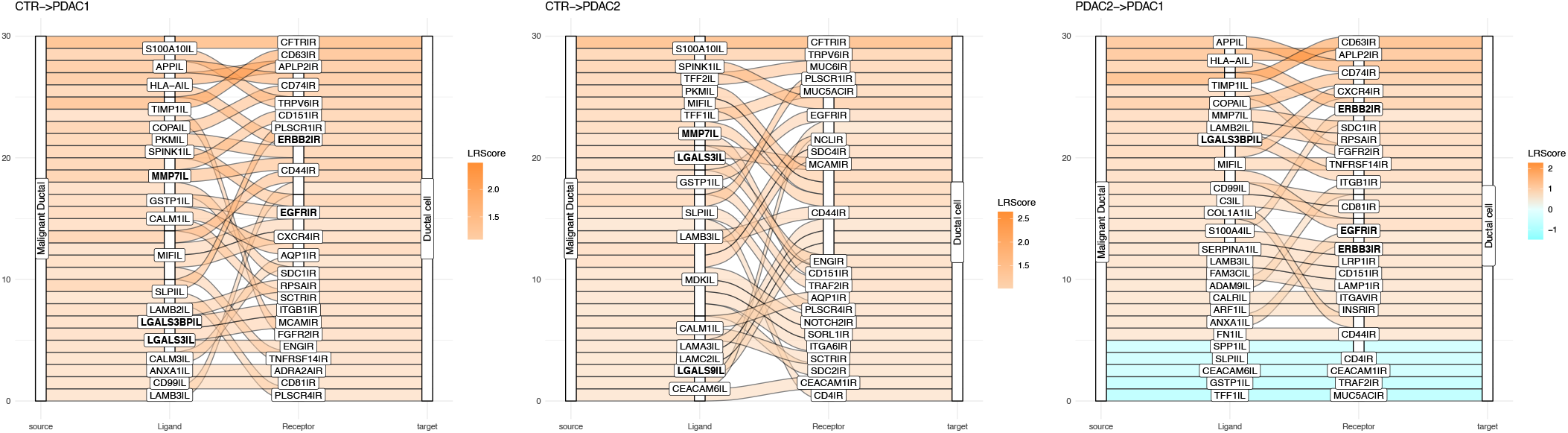
Top 20 ligand-receptor pairs (highest absolute ligand-receptor expression values) between malignant ductal cells towards ductal cells between all three group pairs. We highlight LR pairs discussed in the text.

**Supp Fig 5.**
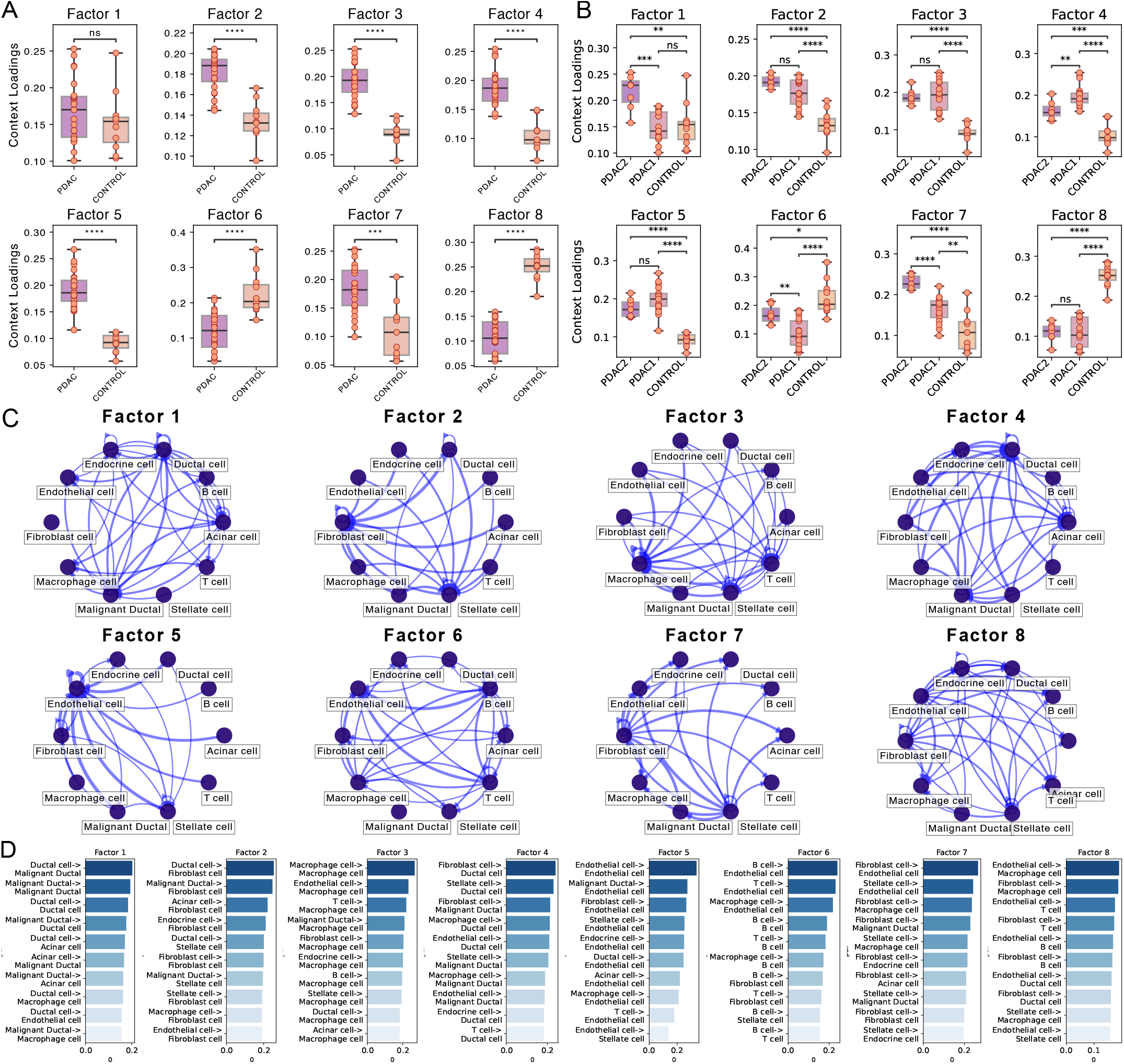
Downstream analysis using Tensor-cell2cell: A) Statistical Comparison between the Context loading using the original disease labels; B) Statistical Comparison between the Context loading using the labels retrieved from scACCorDiON; C) Cell-Cell Communication Networks for each Factor. D) Top 10 highest cell-cell pairs’ context loadings per factor.

